# To draw or not to draw: understanding the temporal organization of drawing behaviour using fractal analyses

**DOI:** 10.1101/2021.08.29.458053

**Authors:** Benjamin Beltzung, Lison Martinet, Andrew J. J. MacIntosh, Xavier Meyer, Jérôme Hosselet, Marie Pelé, Cédric Sueur

## Abstract

Studies on drawing often focused on spatial aspects of the finished products. Here, the drawing behaviour was studied by analysing its intermittent process, between *drawing* (i.e. marking a surface) and *interruption* (i.e. a pause in the marking gesture). To assess how this intermittence develops with age, we collected finger-drawings on a touchscreen by 185 individuals (children and adults). We measured the temporal structure of each drawing sequence to determine its complexity. To do this, we applied temporal fractal estimators to each drawing time series before combining them in a Principal Component Analysis procedure. The youngest children (3 years-old) drew in a more stereotypical way with long-range dependence detected in their alternations between states. Among older children and adults, the complexity of drawing sequences increased showing a less predictable behaviour as their drawings become more detailed and figurative. This study improves our understanding of the temporal aspects of drawing behaviour, and contributes to an objective understanding of its ontogeny.

## 1. Introduction

The expression of artistic behaviour is predominant in *Homo sapiens*, even if the time allocated to such activities tends to decrease with age. A good example of such declining interest across the lifetime can be seen in drawing. Prevalent among children, drawing persists as a recreational activity only among a minority of adults, and as a professional activity among a select few. Drawing develops from internal representativeness (i.e. a drawing representative only from the perspective of the child who produces the drawing) to external representativeness (i.e. a drawing interpretable from the perspective of an independent observer) around the age of 3-4 years (Freeman, 1993; Martinet et al., 2021). Drawing behaviour has been studied in different research fields, especially in psychology, e.g. to characterise its ontogeny (Luquet, 1927; Willats, 2005), its representational development (Adi-Japha et al., 1998; Cherney et al., 2006), the use of colour (Wright and Black, 2013), the comprehension of realism (Jolley et al., 2000) or the influence and role of gender (Cherney et al., 2006; Turgeon, 2008). Such studies have provided critical information about drawing behaviour, despite the fact that the analyses used are often qualitative and subject to biases.

Adi-Japha et al (1998) showed that young children, even if not capable of attributing meanings to the entirety of their drawing, can make sense of broken lines which seem to be more descriptive than curved ones. Although relevant, this method remains subjective because researchers directly question children about what their drawings represent. Moreover, very young children (toddlers) are unable to express themselves with respect to their drawings. Furthermore, the majority of drawing studies focus on spatial measures, whereas the temporal aspects that might also be important in revealing cognitive and decision-making processes remain underexplored. Recent developments in mathematical analyses in animal behaviour might bring new perspectives to the study of such processes.

Indeed, new analytical approaches from statistical physics are now being developed to study individual behaviour, such as fractal analyses. An object is considered fractal when any of its parts examined separately resemble its overall structure at any magnification scale, either in the spatial or temporal domain (Mandelbrot, 1977). A good example of a natural fractal object is Romanesco broccoli, but many other natural systems have fractal structure in the spatial domain, such as river networks (Rinaldo et al., 1993) and human lung architecture (Nelson et al., 1990). Fractal structure is also found in the temporal domain, in processes such as human breathing cycle dynamics (Peng et al., 2002) or human sleep EEG (Weiss et al., 2009). In the behavioural sciences, recent studies have established that fractal analyses of spatial and temporal patterns can lead to a better understanding of animal and human behaviour (MacIntosh, 2014; Rutherford et al., 2003).

On one hand, the study of movement behaviour through spatial fractal analyses have allowed comparisons between observed movement patterns and theoretically optimal foraging patterns (Sims et al., 2008; Viswanathan et al., 1999). We previously applied this approach in examining the efficiency of the drawing trajectory, defined as the correct reading of the drawing with minimal detail in chimpanzees (*Pan troglodytes*) and humans (*Homo sapiens*). Analogous to the trajectory of an animal in its environment, we wanted to know if the efficiency of drawings made on a screen differed between humans and chimpanzees, and with age in humans. Results show that the drawing spatial index was lowest in chimpanzees, increased and reached its maximum between 5-year-old and 10-year-old children, and decreased in adults, whose drawing efficiency was reduced by the addition of details (Martinet et al., 2021).

On the other hand, temporal fractal analyses have shown that an animal’s physiological condition (e.g. health status or degree of physiological stress) affects its sequences of behaviour (Alados, 1996). The concept of fractal time reflects the degree to which current behavioural states depend not only on states immediately preceding them in the sequence but also on those that occur much earlier in the sequence (so-called long-memory processes) (Delignières et al., 2005). Like spatial fractal analyses, temporal fractal analyses also suggest an optimal structure in behaviour time series, within which a normally functioning individual should fall (MacIntosh, 2014). Some environmental pressures or physiological impairments can lead to a loss of complexity in the behavioural sequences of an individual, associated with increases in periodicity or long-range dependence (i.e. greater stereotypy) (MacIntosh et al., 2011; Maria et al., 2004; Meyer et al., 2020). Other factors may in contrast lead to increased complexity, i.e. reduced long-range dependence and increased behavioural stochasticity (Rutherford et al., 2003). To summarise or simplify the concept, certain complexity signatures in behaviour sequences equate to increased stability and/or adaptability, which is the case for some physiological processes as well as sequences of animal behaviour, where it can be an indicator of well-being (Alados, 1996; Maria et al., 2004). By allowing more detailed insight into behavioural sequences, temporal fractal analyses open up new research perspectives for diverse fields of study.

In humans, behavioural sequences are complex and structured in time by multiple factors (e.g., environmental, psychological, physiological), which may limit randomness in the occurrence of behaviour. Several behaviours have been described as fractal, such as patterns of physical activity (Paraschiv-Ionescu et al., 2008) or short-message communication in online communities (Rybski et al., 2009). Considering drawing and handwriting, both pacing and grip force showed a fractal dimension (Fernandes and Chau, 2008). However, the latter study was restricted to an analysis of young adults drawing circles in synchrony with a metronome, so these results cannot be generalised.

In the present study, we sought to improve our understanding of drawing behaviour and its ontogenetic processes by applying temporal fractal analyses to drawing data from humans of all ages. Drawing behaviour is an intermittent process, composed of two main states, including *drawing* (i.e. marking a surface) and *interruption* (i.e. a break in the marking gesture). The temporal structure of each drawing sequence (i.e. sequence memory) can be measured to determine its degree of complexity (MacIntosh et al., 2011). To assess how drawing intermittence develops with age, we collected drawings from 185 individuals of different ages, including children as well as both naive and expert adults. We hypothesized that drawing sequences would be simpler in young children (aged 3 to 5 years), with greater periodicity and long-range dependence in their alternations between states. Indeed, young children who have not yet entered the representative phase of drawing may be more interested in gestures and engaged more in play than in drawing. On the contrary, complexity in drawing sequences may increase among older children and adults, for whom drawings become more and more detailed and representative.

To accomplish our aims, we applied multiple temporal fractal estimators to study long-memory processes in drawing time series. A first challenge was to link these few estimators to drawing behaviour, as we are not aware of any previous studies on this topic. A second challenge was to assess the effectiveness of these fractal estimators, as there is little consensus as to which are most appropriate for a given set of data (Stadnitski, 2012; Stroe-Kunold et al., 2009). Therefore, we first present a robust analytical approach to performing these fractal analyses using our drawing dataset to decrease errors and biases. We then introduce the analysis of complementary indices such as the duration of drawing behaviour in relation to the duration of the drawing session, and the number of swings between drawing and non-drawing states during the drawing session. We discuss our results in the light of biological processes.

## 2. Material and methods

### 2.1 Ethics

We followed the ethical guidelines of our research institutions and obtained ethical approval from the University of Strasbourg Research Ethics Committee (Unistra/CER/2019-11). Drawings were anonymously collected. The contribution of all participants was voluntary and subject to parental consent for children.

### 2.2 Participants

One hundred and forty-four children (67 girls and 77 boys) and forty-one adults (21 women and 20 men) participated in this study (Table 1). We worked with children from kindergarten and primary school in Strasbourg, France. We collected their drawings over two consecutive years (in 2018 for kindergarten children and in 2019 for primary school children). For this reason, 6-year-old children could not be tested in 2019 since they had already participated in the study the year before. Adult participants were between 21 and 60 years old. In addition to a general age effect, we also tested the effect of experience in our adult participants. Among researchers and students at the authors’ research institute, twenty were considered naive (naive adults: 30.8±10.54 years old) as they had never taken drawing lessons and did not draw as a hobby. The other twenty adults were classified as experts and included art school students and professional illustrators (expert adults: 30.4±11.12 years old).

**Table 1.** Groups and gender of human participants.

| Participants groups |  | Gender | Number of participants |
| --- | --- | --- | --- |
|  | 3-year-olds | girls | n=5 |
|  |  | boys | n=15 |
|  | 4-year-olds | girls | n=10 |
|  |  | boys | n=10 |
| Children | 5-year-olds | girls | n=10 |
|  |  | boys | n=10 |
|  | 7-year-olds | girls | n=12 |
|  |  | boys | n=11 |
|  | 8-year-olds | girls | n=9 |
|  |  | boys | n=9 |
|  | 9-year-olds | girls | n=10 |
|  |  | boys | n=11 |
|  | 10-year-olds | girls | n=11 |
|  |  | boys | n=11 |
| Adults | naive | women | n=11 |
|  |  | men | n=10 |
|  | expert | women | n=10 |
|  |  | men | n=10 |

### 2.3 Experimental design

#### 2.3.1. Habituation phase

Children and adults were invited to try the device, a touchscreen tablet (iPad Pro, 13-Inch, version 11.2.2) by drawing on it with their fingers. Participants could use 10 different colours displayed on the bottom of the screen and selected one of them by clicking on it. Adults were tested immediately after habituation but, to avoid overstimulation, children were habituated the day before their respective tests.

#### 2.3.2. Testing phase

Children were tested individually at school, either in their own classrooms (for 3-year-olds) or in the staffroom (for older children). During the test, the experimenter (LM or MP) stayed with the child but kept their distance so as not to influence their behaviour. Adults participated individually, either alone in a room at the research institute (for naive subjects) or at the art school (for experts). We used a video camera to record the hand movements of each participant while drawing, in case we needed to control for any contextual issues arising during the session (e.g. disruption of the drawing session, unintentional points or lines made by the experimenter at the end of the session, *etc*.). No time limits were imposed during the study.

Participants were tested under two conditions to assess potential differences between a non-specific task (*free* drawing) and a specific task (drawing a *self-portrait*). Under the *free condition*, subjects were asked to draw whatever they wanted, with no further instructions. Under the *self-portrait condition*, subjects were asked to draw themselves. In each participant category, we randomly assigned subjects such that half began the test with the *free condition* instructions while the other half began with the *self-portrait condition*. To avoid overstimulation and a lack of concentration, none of the children participated in both conditions on the same day.

#### 2.3.3. Data collection

Three hundred and sixty-nine drawings were collected (one drawing by a naive adult was deleted by mistake). Since there were no imposed time limits, so as to not constrain the creativity of each participant, the drawing duration was different for each person and ranged from 17 seconds to 923 seconds (mean ± s.d. = 250.5± 189.2). When the individual was drawing, a triplet (x,y,t) was recorded every 17 milliseconds on average (resolution of point recording), where (x, y) is the position of the finger on the touchscreen and t the time. For two consecutive points (x_i_,y_i_,t_i_) and (x_i+1_,y_i+1_,t_i+1_), the time interval in milliseconds is t_i+1_ - t_i_. If t_i+1_ - t_i_ < 100, we considered it drawing time. If t_i+1_ - t_i_ > 100, we considered it non-drawing time; i.e., the individual must raise their finger long enough for this stop to be voluntary (Tanaka et al., 2003). This succession of time intervals was then transformed into a binary series of -1 (not drawing) and 1 (drawing), where the duration between two consecutive points was set at 1 millisecond. Note that analyses with other resolutions (10 milliseconds and 17 milliseconds) were performed for comparison and gave similar results (see Supplementary Materials Figure 1). The resultant time series can be represented as a barcode as shown in Figure 1. The width of each black band corresponds to a drawing duration, while a non-drawing duration is equivalent to the width of a white band. Each video was watched several times to remove moments during which participants were distracted by something else (Longstaff and Heath, 1999) (simulations showed that this did not mathematically impact the results, see Supplementary Materials Box1). This happened a few times at school during drawing sessions with children. This concerns only 22 drawings in our final analyses (i.e. 6.4% of the dataset) where 1 to maximum 2 durations were removed, representing 12.2±7.1% of the final sequence (these were generally times exceeding 15 seconds on average and caused by a child entering the room, the bell for recess ringing, *etc*).

**Figure 1.**
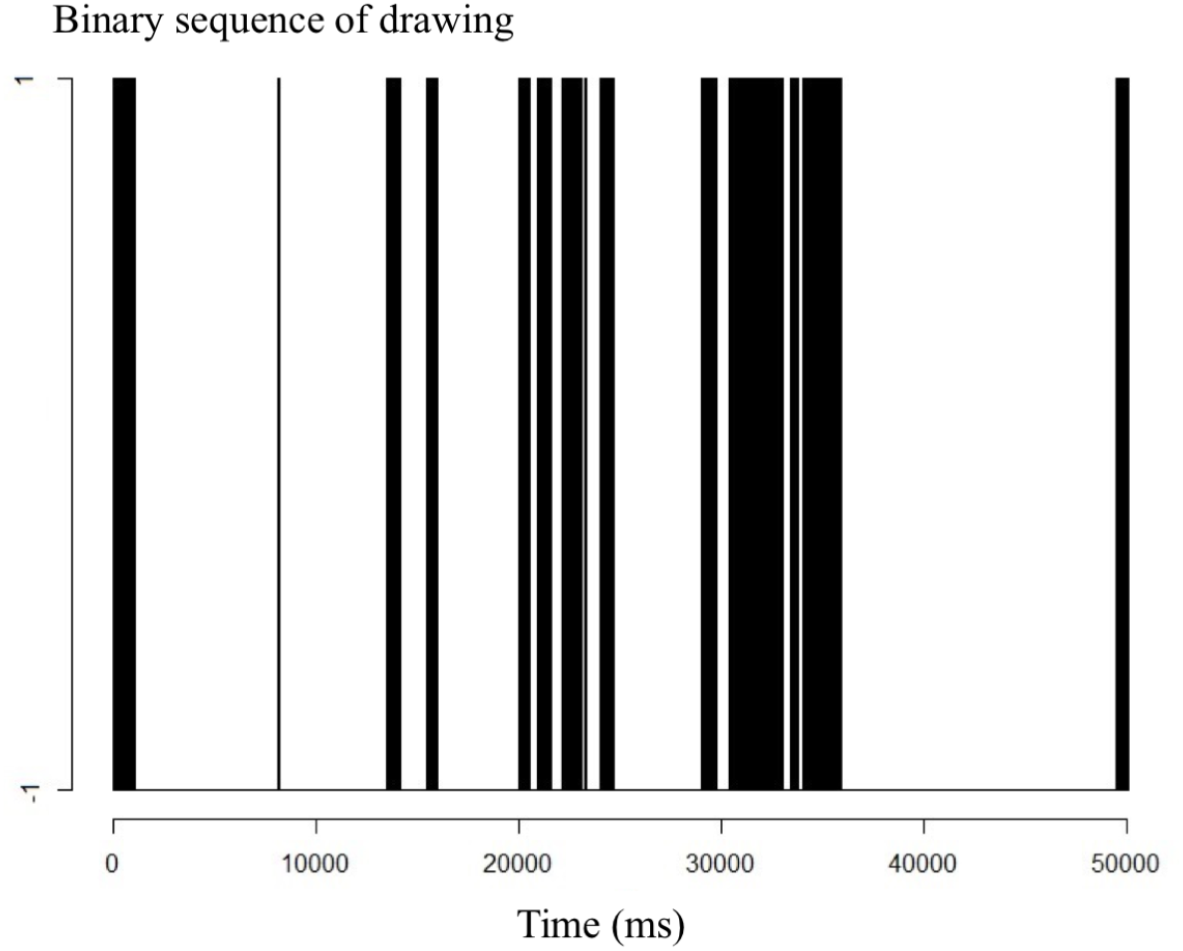
Example of a binary sequence of intermittent drawing, denoted 1 for drawing and -1 for non-drawing behaviour. Black bars reflect the durations of the drawing state while white bars reflect the durations of the non-drawing state.

### 2.4 Analyses

#### 2.4.1. Type of signal

Fractal characteristics or long memory processes can be measured via different algorithms, each having its own statistical parameter. Here, the difficulty arises from the fact that, for each parameter, numerous estimators have been defined, but the effectiveness of each is still debated in the literature (Stadnitski, 2012; Stadnytska et al., 2010). Studies often focus on one or few estimators without a rigorous reason for comparing them. As a consequence, there is no simple and systematic way to estimate the fractal process, which often results in errors or misleading conclusions (Karagiannis et al., 2006).

The most widespread way to assess and quantify long memory processes in temporal sequences is estimation of the Hurst exponent (*H*). This exponent is a measure of the correlation among signal components in a time series (Cannon et al., 1997; Stroe-Kunold et al., 2009) Within this framework, behavioural sequences can reflect three mutually-exclusive scenarios: (1) *persistence* (*H*>0.5) occurs when positive long-range autocorrelation exists, such that blocks of certain behaviours (e.g. drawing) are likely to be followed by blocks of similar duration in succession; (2) *anti-persistence* (*H*<0.5) occurs when negative long-range autocorrelation exists, such that blocks of behaviour are likely to be followed by blocks of divergent duration; (3) *white noise* (*H*=0.5) occurs when no sequence memory exists, i.e., the sequence is random or contains only short-range autocorrelation (Delignières et al., 2005).

Methods for *H* estimation differ depending on the signal class of the original sequence, which can be either *Fractional Gaussian noise* (fGn) or *fractional Brownian motion* (fBm) (Mandelbrot and Van Ness, 1968). fGn is stationary with constant variance and mean whereas fBm is nonstationary, even if both signals are theoretically linked: differencing fBm produces fGn and integrating fGn creates fBm (Stadnytska et al., 2010). The same original sequence expressed as one or the other signal class will be characterized by the same Hurst exponent (Figure 2). However, before estimating a behaviour sequence’s *H* exponent, it is essential to first define its signal class, which can be done through different methods (Cannon et al., 1997). The so-called ‘temporal’ methods are those which do not require prior transformation of the data and identify statistical dependence in elements of the time series (Eke et al., 2002). Frequency-based methods, on the other hand, are based on transformation of the time series, for example by considering the periodogram (i.e. the spectral density of a signal). In this article both types of methods are used with the aim of comparison and to combine the different approaches for a more robust investigation.

**Figure 2.**
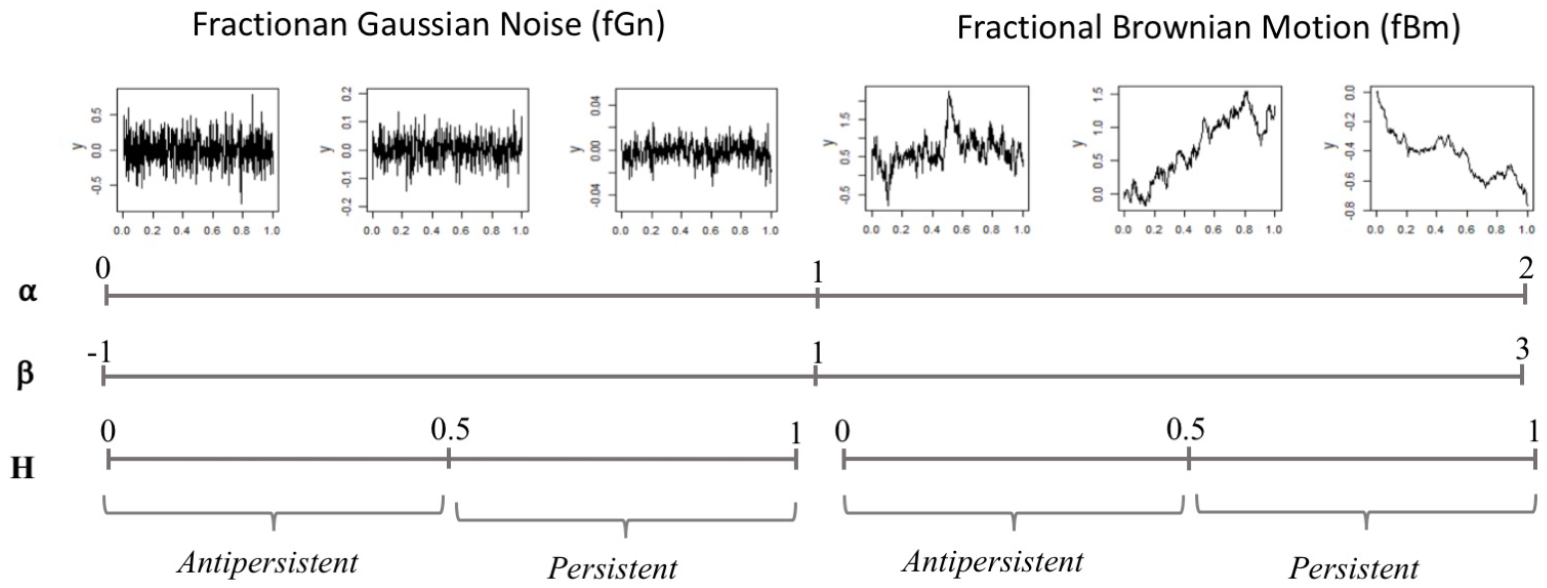
Illustration of the fGn/fBm continuum. Values of the scaling exponent *α* (from detrended fluctuation analysis: DFA) and the index *β* (from power spectral analysis) are depicted in relation to the Hurst exponent, *H*. This representation is largely inspired by (Marmelat et al., 2012).

#### 2.4.2. Temporal methods

##### Detrended Fluctuation Analysis (DFA)

To investigate long-memory processes in the sequential distribution of drawing and non-drawing durations, we employed Detrended Fluctuation Analysis (DFA) (Peng et al., 1995) which is among the most used to study binary sequences of animal behaviour (MacIntosh et al., 2013; Meyer et al., 2020; Rutherford et al., 2003). It is also a robust estimator of the Hurst exponent (Cannon et al., 1997; Eke et al., 2002). DFA calculates a scaling exponent (*α*_DFA_) corresponding to the slope of the line on a log-log plot of the average fluctuation at each box size (given by the three steps, from equation 1 to equation 3 below). A lower *α*_DFA_ reflects greater stochasticity and reduced long range memory (Figure 2) (Meyer et al., 2017). If a linear relation exists, it indicates the presence of scale invariance.

To summarize the application of DFA to our data, a binary sequence of a drawing (Figure 3a) is cumulatively summed such that

**Figure 3.**
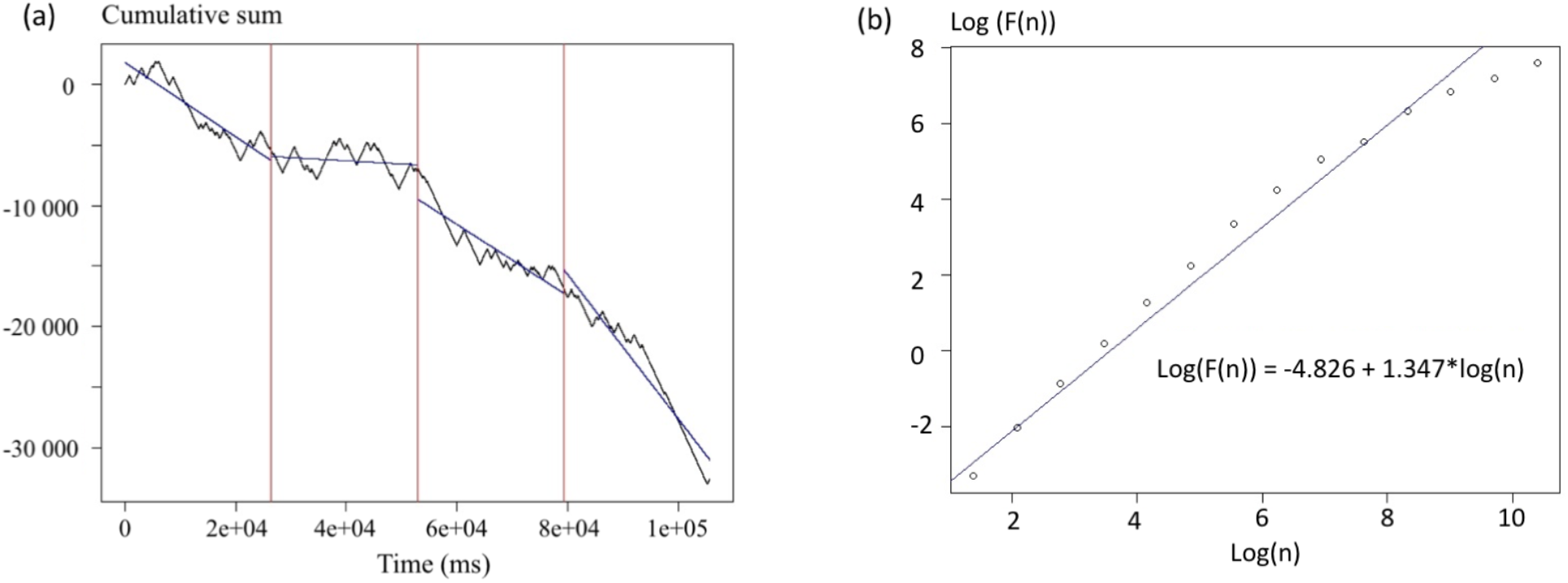
(a) Integration of a drawing sequence. Vertical lines give an example of dividing the sequence into N/n = 4 boxes. Lines in each box correspond to the polynomial regression. (b) Bi-logarithmic plot of the statistic *F(n)* against the length of the time intervals *n*. The regression line allows calculation of the scaling exponent *α*, which is its slope (1.347 in this case).

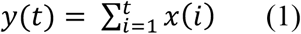

Where *y(t)* is the cumulative sum and *x(i)* is the binary sequence at each time step (1ms intervals).

The cumulative sum is then divided into non-overlapping boxes of length *n* (Figure 3a) and a least-squares regression line is fit to each box *y*_*n*_*(t)* to remove local trend and it is repeated over all box sizes, then the fluctuation is calculated as follows (Figure 3b)

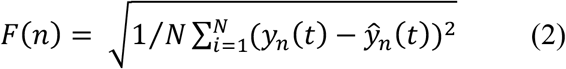

Where *ŷ*_*n*_(*t*) is the regression estimate for *y*_*n*_ (*t*) at each box size *n* and *F(n)* is the fluctuation of the modified root-mean-square equation across all scales available (2^2^, 2^3^, 2^4^… 2^m^) where *m* is the largest scale examined, such as 2^m^< N/2 (rounded down to the nearest whole number) (Constantine and Percival, 2007; Stadnytska et al., 2010; Stroe-Kunold et al., 2009). We then obtained the following relationship

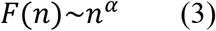

This method makes it possible to know whether the starting signal is a fBm or a fGn, since it can be used on both (Seuront, 2009; Stadnytska et al., 2010; Stroe-Kunold et al., 2009). The *α* exponent can be interpreted as follows:

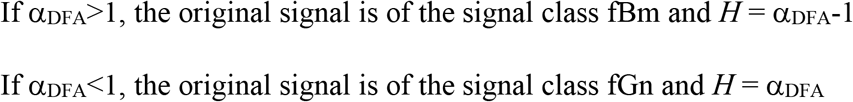

*α*_DFA_ in the range of 0.5–1 and 1.5–2 reflects persistence while *α*_DFA_ in the range of 0–0.5 and 1–1.5 reflects anti-persistence for fGn and fBm, respectively (Figure 2). Values of 0.5 and 1.5 reflect Gaussian (white) noise and Brownian motion, respectively.

##### Hurst Absolute Value (HAV)

The HAV method is similar to DFA but does not integrate the time series before analyses so that *H* is calculated from the original binary sequences of drawing and non-drawing durations. We included the HAV method because it is able to capture the self-similarity parameter in time series data where DFA fails to do so (Mercik et al., 2003). This method only works on fGn, so any application of HAV on fBm signals must use their increments (i.e. the corresponding fGn). A time series of class fGn of length *N* is divided into smaller boxes of length *n*, denoted as *x*^*(n)*^, and the first absolute moment is obtained as follows:

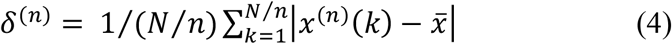

This is reiterated for the different window size *n* with the variance *δ* varying as follows:

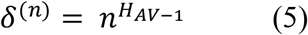

Where H_AV_ is the scaling exponent.

##### Scaled Windowed Variance (SWV)

Since we only had fBm series, the SWV method was selected as it a good estimator for this type of signal (Delignieres et al., 2006). The fBm series *x*(*t*) is divided into non-overlapping boxes of length *n*. In each box, a bridge detrending method is used to remove the trend, this is what is recommended for series of more than 2^12^ points (Cannon et al., 1997), which is our case here. Then the standard deviation (SD) is calculated in each box such that

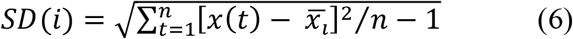

With 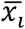 being the average in the box *i*. Then, the average standard deviation 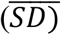 of all boxes of length *n* is calculated such that

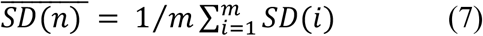

Where *m* is the number of boxes of length *n*.

This step is then repeated over all possible box lengths and 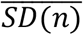 is related to *n* by a power law

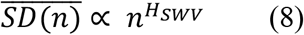

The 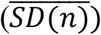 is plotted against log(n) and the slope of the regression is the estimated Hurst coefficient H_SWV_ (Cannon et al., 1997; Delignières et al., 2005).

#### 2.4.3. Frequency-based method

##### Power Spectral Density analyses (PSD)

Like DFA, this method is widely used to determine the signal class of time series data. It is included here for comparison, because of the widespread use of PSD in standard time series analysis, and the added diversity it affords us as an index based on frequency. In the frequency domain, the fractal character is expressed through the following power law:

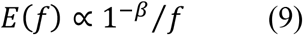

where *E(f)* is the squared amplitude for the corresponding *f* frequency. The *β* exponent is obtained by calculating the negative slope (*-β*) of the regression between *log(E(f))* and *log(f)* (Figure 4). *β* values range between -1 and 3, with *β* between -1 and 1 reflecting fGn and *β* between 1 and 3 reflecting fBm (Delignières et al., 2005; Eke et al., 2000). *β* is linked to *H* such that:

**Figure 4.**
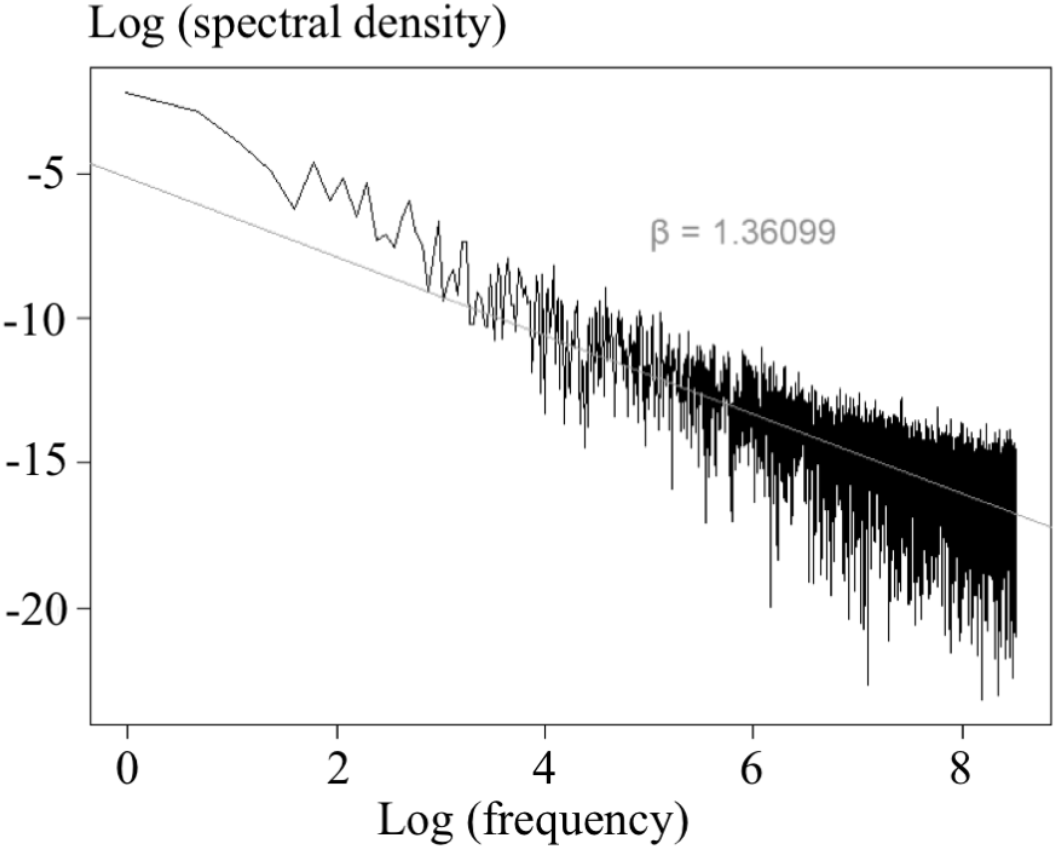
Bi-logarithmic power spectrum of a time series with regression slope fitted to obtain the *β* index.

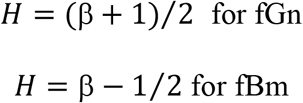

In the present study, the high frequencies (1/8<f<1/2) are excluded. This method provides more reliable estimates of *β* and is known as ^low^PSD (Stadnitski, 2012).

#### 2.4.4 Methods implementation

We performed DFA using the package *fractal* (Constantine and Percival, 2007) in R. For the Power Spectral Density, we based our analyses on the script found in Stadnitski et al. (2012). As the Scaled Windowed Variance and Hurst Absolute Value methods are not currently implemented in R, we coded and tested our own algorithms through Monte-Carlo simulation. We simulated, using the *somebm* package (Huang, 2013), fBm of different lengths and known *H*. In these simulations, the lengths of the series were defined as powers of 2, from 2^11^(= 2,048 points) to 2^15^ (= 32,768 points). For each possible length, 100 fBm were generated with *H* = 0.2, *H* = 0.4, *H* = 0.6 and *H* = 0.8. More than 2000 series were generated in total and the errors,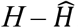, are shown in the Supplementary Materials (Figures 2 and 3). The drawing sequences obtained in this study were much longer (mean ± s.d. = 240,832ms±185,561ms) than these simulated series and therefore will not be a limiting factor in the estimates. Since the HAV method only works on fGn, the simulated series used are those of the respective fBm increments. These simulations have shown that the larger the size of the series, the closer the errors are to 0, proving the efficiency of these analyses.

#### 2.4.5 H estimation

The Hurst coefficient *H* was estimated using the four methods described above. The binary time series studied all being fBm, it was necessary to consider series consisting of the increments of these fBm (i.e. the corresponding fGn) to be able to apply certain methods such as HAV. As mentioned earlier, each of these methods does not always directly estimate the Hurst coefficient, *H*, but the latter’s relationship with the calculated exponent is relatively simple (Figure 2). Once the four methods were applied to each series, a Pearson correlation matrix was calculated.

#### 2.4.6 Additional temporal indices

Both of the indices described below were extracted from the complete binary sequences.

##### Proportion of drawing in the sequence

We first calculated the number of 1’s in the sequence. Since each binary sequence had a different length, we divided the sum of 1 by the total sequence length to come up with a drawing proportion for comparison across sequences.

##### Rate of state changes

The number of state changes corresponds to the number of times a participant changed their behaviour between drawing and interruption during the drawing session. For each sequence, the number of state changes was divided by the total sequence length and multiplied by 1000 to obtain a rate of change per second.

#### 2.4.7 Statistical analysis

Statistical analyses were conducted in R (version 3.6.2). Since we used four methods to estimate *H*, we obtained 4 estimated values of this index for each time series to use in our statistical analyses. Some previous studies have advised averaging the different estimates, but the choice of which to use and the overall number of estimators used has varied (Eke et al., 2000; Seuront, 2009). The four estimates of *H* were found to be highly correlated with each other. Considering that averaging variables whose variance may be different can remove information, we chose to perform a PCA. In our case, the method was used to reduce the number of estimators used into one that retained as much of the information as possible. The new resultant variable was thus constructed as a linear combination of the original variables, allowing for a synthesis of our variables into a single index (Berni et al., 2011). After using the Kaiser-Guttman criterion to select the number of axes, only values of the first principal component from the PCA – a proxy for the Hurst exponent that we term the ‘*Hurst axis*’ – were used and set as a response variable in our statistical models. In our case, results obtained following this methodology with those resulting from averaging the estimates were equivalent (see Results). We decided to retain the PCA, as it makes it possible to create a new variable (i.e. an index) which is an optimally weighted combination of correlated estimators of *H*.

A correlation of -0.43 was observed between the sequence length and the Hurst axis. Indeed, there were considerable differences in the lengths of our drawing sequences, which averaged approximately 4 minutes and ranged between approximately 17 seconds and 15 minutes (mean ± s.d. =240832±185561ms; range=16848-908250ms). We were not interested in this correlation but needed to account for it when testing relationships between *H* and our other variables of interest, such as the *gender*, the *group* (age) or the *condition* under which the drawing was made. For this reason, the four estimated coefficients of *H* were recalculated on the first 50,000 points (i.e. the first 50 seconds of the drawing), 100,000 points, 150,000 points and 200,000 points. DFA coefficients based on the first 50,000 points in the sequence were correlated with those based on these other three lengths (Supplementary materials, Figure 4), as well as with those from the whole sequence at more than 54%, meaning that the information contained in the first 50,000 points is a good threshold compared to that contained in the entire sequence. Therefore, we analysed only the first 50,000 points of each sequence and excluded all sequences less than 50,000 points in length (6.5% of the drawings collected).

Given the above criteria, we analysed 346 drawings out of the initial 369 we obtained. After recalculating the indices on the first 50,000 points for each sequence, the PCA procedure was redone. We then determined whether *group, gender* or *condition* were associated with variation of the “Hurst axis” by constructing a Generalized Linear Mixed Model with Gaussian error structure (*nmle* package (Pinheiro et al., 2006)). Since each participant produced two drawings, *individual identity* was added as a random factor. Residual normality was graphically verified. The normality of the residuals of the random effect was graphically validated for each group. Since heteroscedasticity was detected across groups in the original model, we added a covariance structure (*VarIdent*, adapted to the categorical variables) to allow the variance of the residuals to change according to group. The full model included all possible variables and first-order interactions between *group, gender*, and *condition*. We proceeded to model selection using a dredge function based on the lowest Aikake’s Information Criterion (package *MuMIn*; ΔAIC > 2 (Barton, 2009)). Paired comparisons were made using the *lsmeans* package to compare different age groups in pairs.

Concerning the two additional metrics, the proportion of drawing in the sequence and the rate of state changes, measures have been done on the binary sequences. For the number of states changes, we normalized the data using a Box-Cox transformation and ran a Linear Mixed Model (package *nlme*) containing the variables *group, gender* and *condition* with *individual identity* added as a random factor. Again, model selection was carried out using the dredge function (package *MuMIn*) and we chose the model with the lowest AIC. Residual normality was graphically verified, but since heteroscedasticity was detected, we added a covariance structure to allow for a difference in the residuals variance between groups (*VarIdent*). For the proportion of drawing, we used a Linear Mixed Model (package *nlme*) too. The conditions of application (residual normality and homogeneity of variance) were graphically verified. The alpha level for all statistical analyses was set at 0.05.

## 3. Results

### 3.1 Type of signal

Examination of *α*through DFA(mean ± sd = 1.419 ± 0.0490) shows that the original binary sequences were characteristic of fBm. This result was confirmed by the ^low^PSD method which estimates of *β* were greater than 1 (mean ± sd = 1.927 ± 0.0472).

### 3.2 Estimates of H

The Hurst exponent was estimated with the four methods (Supplementary Materials Figure 5). The binary time series studied being all fBm, we considered series resulting from the increments of these fBm. Each of the methods is positively correlated to the others, an expected result since they estimate the same coefficient (Figure 5).

**Figure 5.**
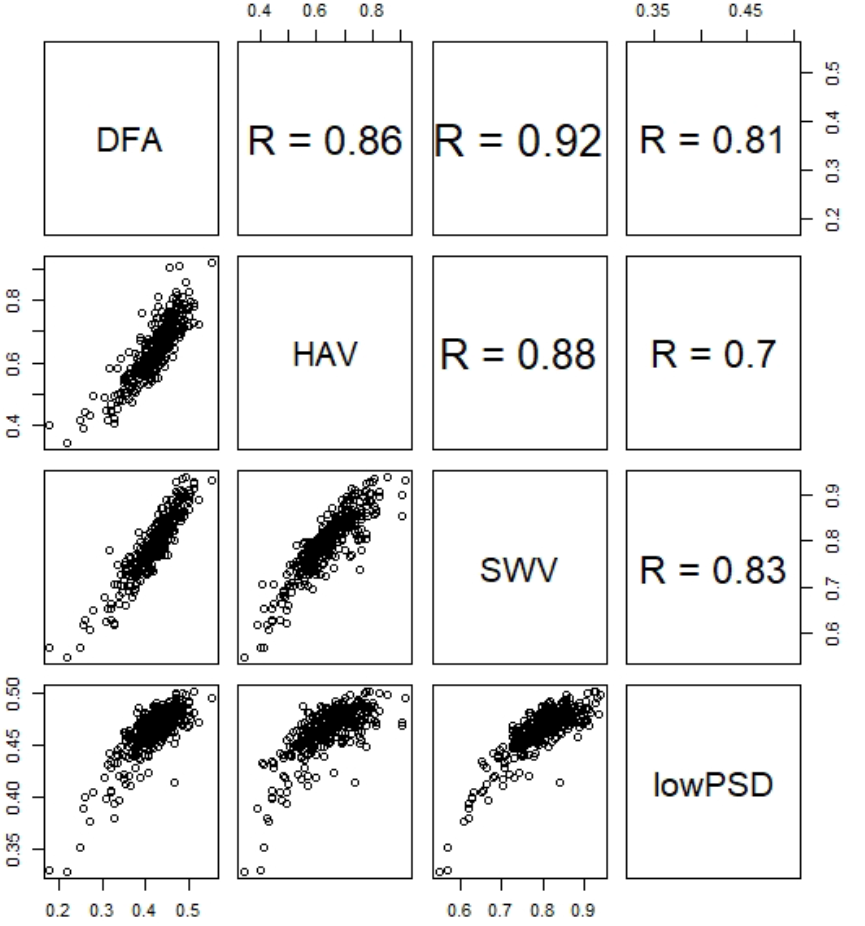
Pearson correlation matrix indicated the correlation values between the four methods used to estimate the Hurst exponent H.

### 3.3 Combination of the estimators through a Principal Component Analysis (PCA)

Since the PCA was standardized, only the axes whose inertia was strictly greater than 1 were kept, which was equivalent to keeping only axis 1in our data set, explaining 87.68% of the variance (Correlation circle available in Supplementary Materials Figure 6). All *H* estimators loaded positively into the first principal component, i.e. the Hurst axis (the loadings for DFA, HAV, SWV, and ^low^PSD were 0.513, 0.491, 0.518, and 0.475, respectively).

**Figure 6.**
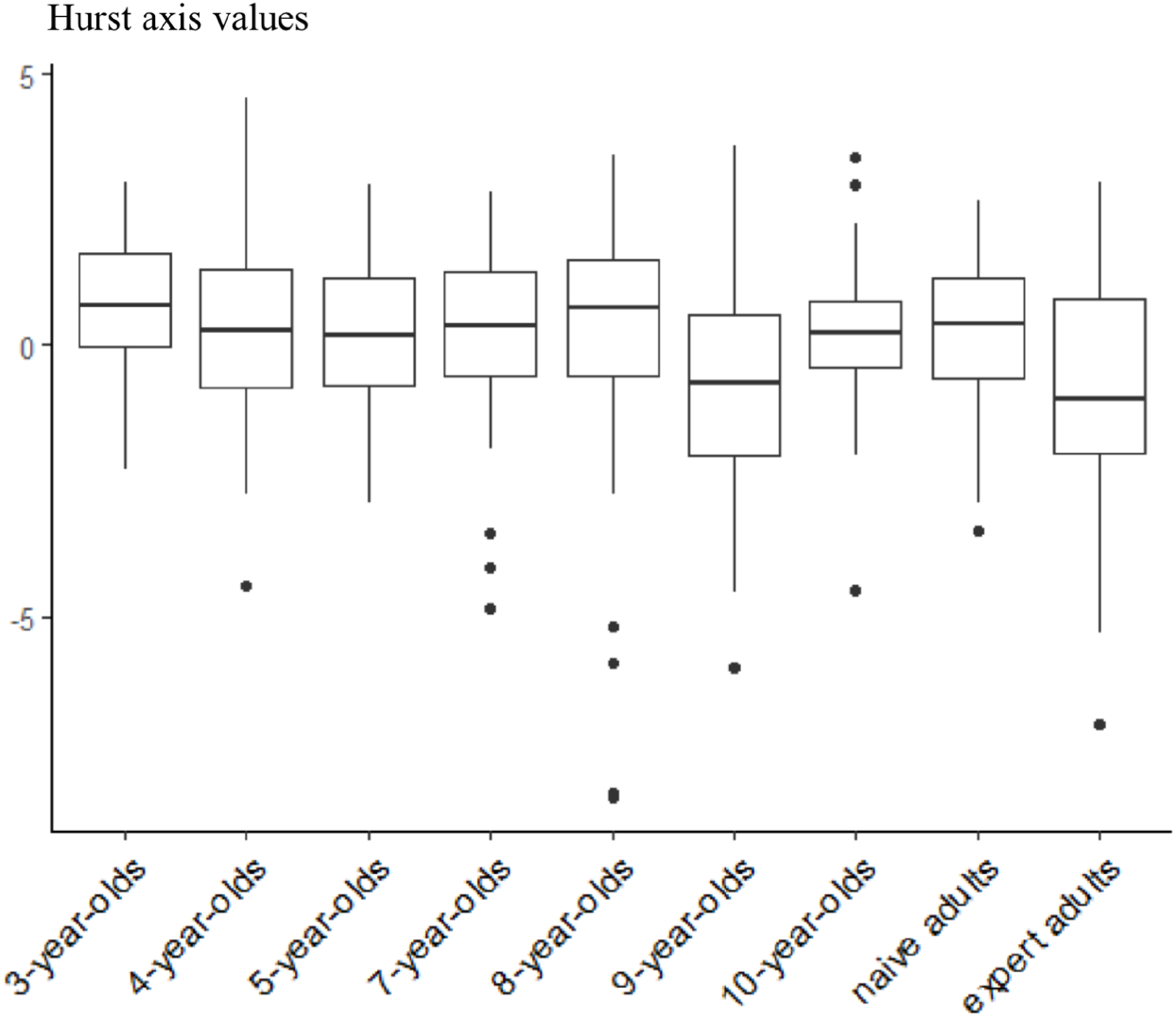
Boxplots of the Hurst axis values for each group. Each boxplot depicts the median (bold bar), 25-75% quartiles (box) and outliers (points).

### 3.4 Variation in the Hurst axis according to the group

The selected model was the one that contained only the variable *group* (*df* = 8, χ^2^= 21.434, p = 0.006; Figure 6). Paired comparisons indicated that two significant differences emerged: 3-year-olds had a higher value along the Hurst axis than novice (p = 0.0085, t = 3.706) and expert (p = 0.0148, t = 3.540) adults. Neither gender nor condition was associated with variation in the Hurt axis.

Furthermore, values of the Hurst axis were highly correlated with the average value across our 4 estimators (*r* = 0.993), and the same statistical model, containing only group as an independent variable, best explained variation in this averaged metric (Supplementary Materials Table 1).

### 3.5 Additional temporal indices

#### 3.5.1 Proportion of drawing in the sequence

The selection of models led us to choose the model containing the variables *group* (*df* = 8, χ^2^ = 65.559, p < 0.0001) and *condition* (*df* = 1, χ^2^ = 13.042, p = 0.0003). Naive adults showed a proportion of drawing significantly lower than that of all other participants (p < 0.005, t < -3.845; Figure 7). In addition, the proportion of time spent drawing as higher in the *free* condition compared to the *self-portrait* condition (p = 0.0004, t = 3.611; Figure 7).

**Figure 7.**
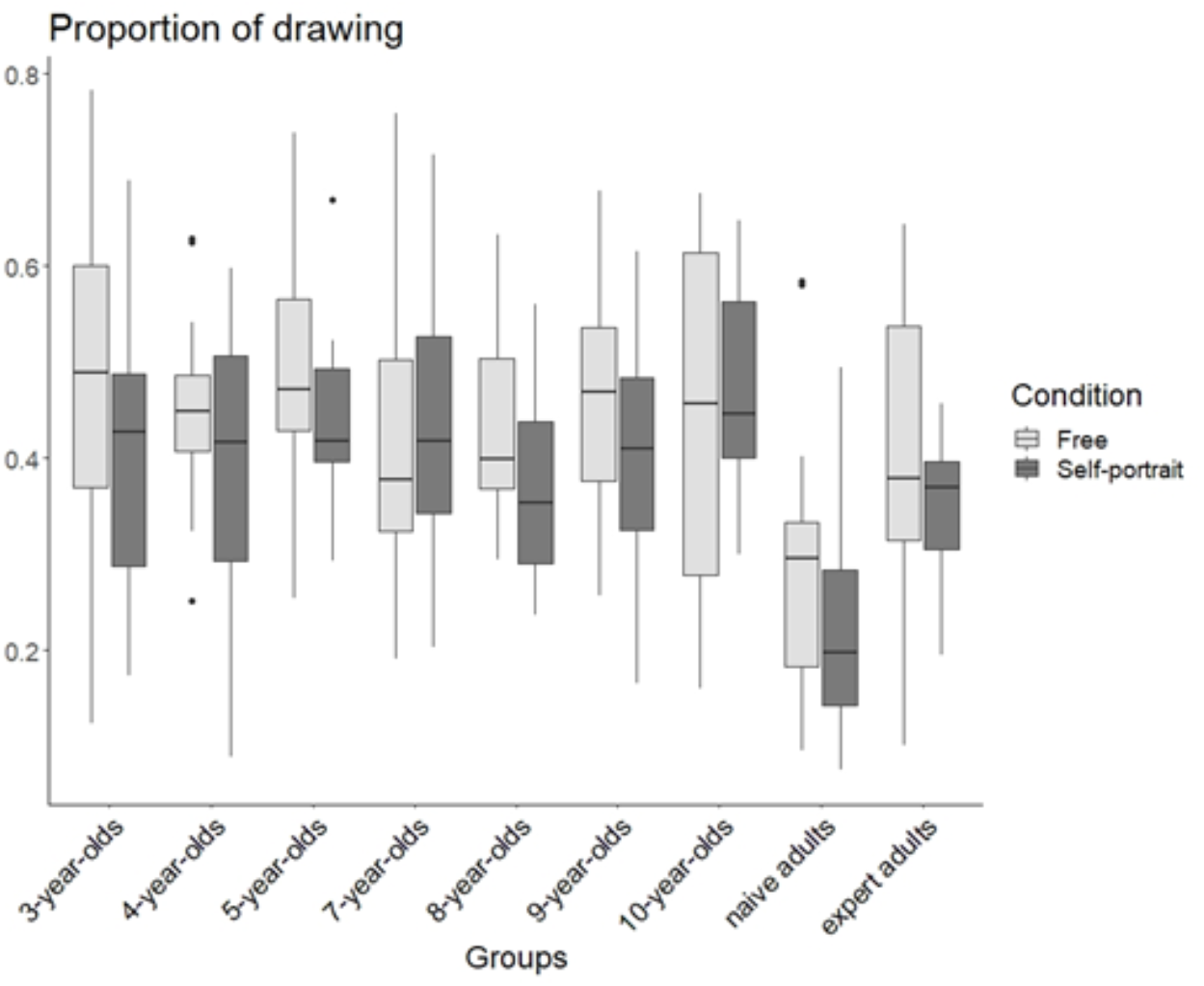
Boxplot of the proportion of drawing for each group and for both conditions. Each boxplot depicts the median (bold bar), 25-75% quartiles (box) and outliers (points).

#### 3.5.2 Rate of state changes

The selected model was the one containing only the variable *group* (*df* = 8, χ^2^ = 32.607, p <0.0001). The 3-years-old children alternate significantly less between drawing and interrupting behaviours than 5-year-olds (p = 0.0310, t = -3.303), 8-year-olds (p = 0.0262, t = -3.359) and adults, both naive (p = 0.0011, t = -4.261) and expert (p = 0.0009, t = -4.303) (Figure 8). No other significant effects, gender or condition, were found.

**Figure 8.**
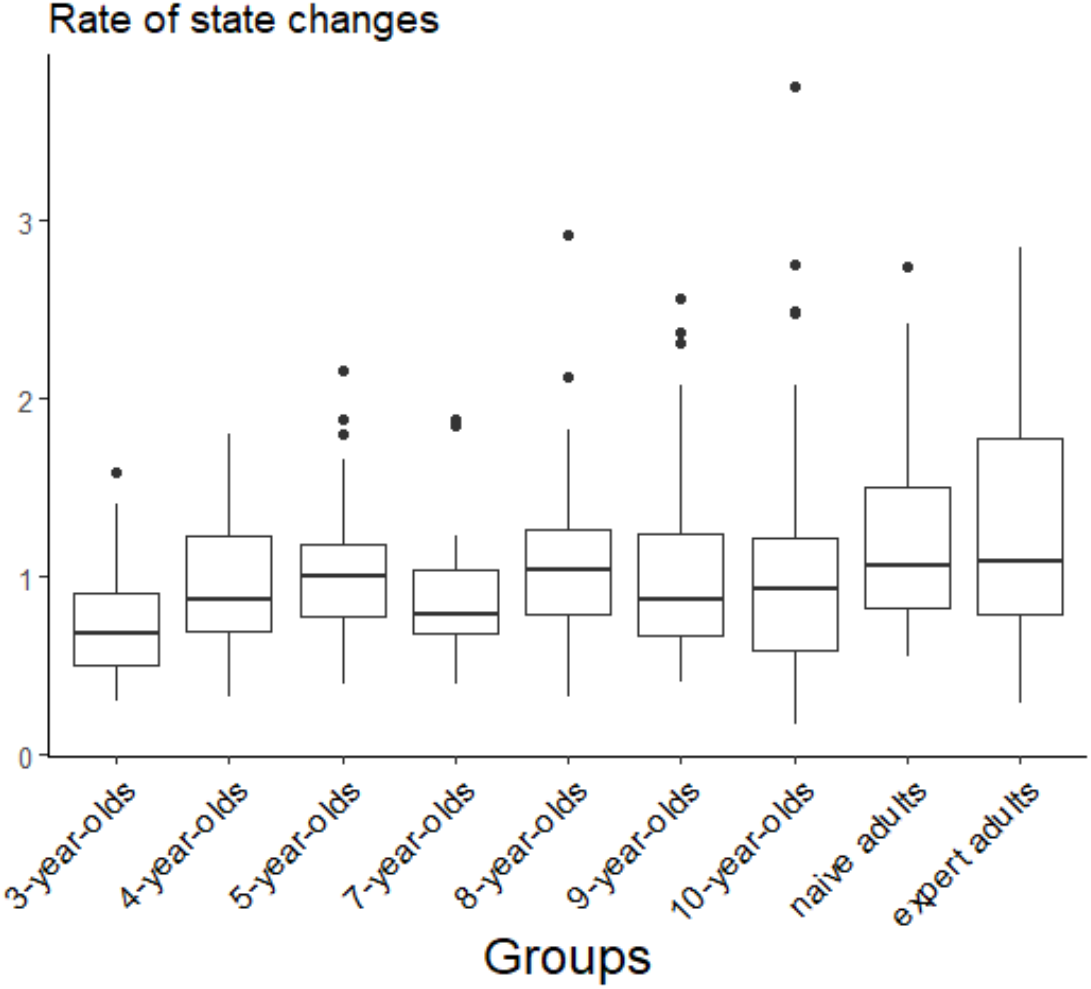
Boxplot of the rate of state changes (number of state changes per second) for each group. Each boxplot depicts the median (bold bar), 25-75% quartiles (box) and outliers (points).

## 4. Discussion

With this study, we wanted to know whether temporal fractal analysis could provide insight into the study of drawing behaviour, as has been done in other fields (MacIntosh, 2014) but not yet – to our knowledge – to understand intermittence in drawing. Specifically, we tested whether the temporal dynamics of drawing varies according to the age, gender or instruction under which the drawing was carried out. In a previous study applying spatial fractal analyses in the field of drawing behaviour, we found that the efficiency of the drawing trajectory, defined as the correct reading of the drawing with minimal detail, increased during childhood and reached its maximum in children between 5 and 10 years of age, before decreasing in adults due to the addition of greater detail (Martinet et al., 2021). Studying temporal aspects should allow us to further understand the ontogeny of drawing behaviour.

We found a difference between the youngest (3-year-old children) and oldest (adults) participants, meaning that these two groups do not draw with the same patterns of alternation between drawing and non-drawing states. In other words, the process of drawing shows different degrees of temporal complexity. Young children showed the highest values of the Hurst axis, suggesting that they exhibit more stereotypical and therefore less complex temporal patterns in their drawing behaviour. On the contrary, temporal sequences of drawings were found to be more complex, meaning less predictable, in both naive and expert adults. In relation to this, the rates of state change we observed demonstrate that 3-year-olds performed significantly fewer alternations between drawing and drawing interruptions than all adults, 8-year-olds and 5-year-olds. Said differently, the drawings of the youngest children are composed of fewer strokes per unit of time spent on the activity.

In general, previous work has shown that 3-year-old children draw for shorter periods than older participants, get bored faster and may be more motivated by the simple motor pleasure of using the tablet rather than recognizing it as a real drawing support tool (Martinet et al., 2021). Many young children first tried each available colour, one by one, which might have induced a certain stereotypy in alternations between drawing and interruption, leading to high Hurst axis values. Their drawings were comprised of what could be called scribbles, as they were not figurative, not representative, at least to the eye of an external observer (Martinet et al., 2021).

However, 3-year-old children did not stand out in terms of the proportion of time spent drawing during a session. Indeed, all participants spent a greater proportion of their time drawing in the *free* condition than in the *self-portrait* condition, and we know from a previous study that the durations of the entire drawing sessions were also longer in the *free* condition compared to the *self-portrait* condition (Martinet et al., 2021). The present study confirms that the addition of instructions limits the proportion of actual drawing time during a session by requiring more reflection time.

Regardless of the condition, the proportion of time spent drawing during a session was significantly lower in naive adults. Naive adults, more than any other group, expressed feelings of being judged and apprehension toward doing wrong, and expressed explicitly that they did not know how to draw. Despite these apprehensions and their alleged impacts on performance, the *H* estimates characterizing the drawing behaviour of naive adults was not different from those of experts. Whether this suggests that different mechanisms can lead to similar fractal patterns in drawing, or that such patterns are not sensitive to subjective experiences during drawing, cannot be determined at this time. However, this does highlight that the differences between young children and adults may not depend strongly on experience or skill but may instead reflect more fundamental ontogenetic processes.

Given that the majority of drawings made by 3-year-olds did not exhibit external representativeness, one possible explanation for this difference of complexity with age could be a desire for figuration. Indeed, in adults the process of drawing is intentional and may lead to greater stochasticity in the intermittences between drawing and non-drawing states, due to thought processes and/or tendencies toward representativeness. When an individual produces a figurative drawing, recognizable by an observer, the intentionality of his acts is obvious. Concerning abstract drawings, non-figuration does not necessarily mean absence of intention. The probable role of drawing instructions can then be evoked. This last consideration leads us to go further in future drawing analyses, asking adults to draw abstract or figurative drawings for comparison with children’s’ scribbles. As the temporal fractal index is a measure of the temporal complexity of the drawing behavioural sequence, if the two measures of H are different between the abstract drawings made by young children and those made by adults, this would mean that the complexity assessed by the H index could be interpreted as a measure of intentionality more than just figuration. Additional studies would be needed to confirm this.

In the present study, drawing instructions (*free* and *self-portrait*) had no effect on drawing intermittence. Extrapolating from the previous discussion, inviting participants to produce archetypes, more stereotyped drawings of common objects such as a house or a flower, might produce a gradient of temporal patterns depending on the complexity of the task. Perhaps asking subjects to draw a specific object would lead to homogenization of temporal patterns across participants. Asking participants to reproduce a photograph – which would reduce the role of creativity and therefore minimize reflection time but not necessarily simplify the drawing task – may lead to further variation, and potentially reduced variability between subjects. The latter would allow measuring the variation in drawing complexity while removing cultural and normative aspects. In this way, it would be possible to reach stronger conclusions about the influence of a directive on drawing behaviour.

Concerning differences in drawing behaviour between individuals, gender can be an influential factor. Previous results have shown such differences in the fields of drawing and writing, particularly with regards to colour utilization, where girls show a more extensive use of colour than boys (Martinet et al., 2021; Turgeon, 2008; Wright and Black, 2013). However, we found no evidence to suggest that drawing intermittence differs between girls/women and boys/men.

Though we observed clear differences in fractal patterns of drawing intermittence between age groups, there remain limitations to the study. For example, difficulty arises from the fact that multiple methods exist to estimate the Hurst exponent, which often leads to conflicting results (Karagiannis et al., 2006). It remains challenging to determine which estimator best suits the type of data being analysed, and this is exacerbated by the fact that estimations themselves vary according to the method used even in simulated time-series with *a priori* seeded Hurst values (Karagiannis et al., 2006). In our study, individuals with high (or low) H according to one method generally exhibited similarly high or low H with other methods, but the variations between groups differed from method to method. Indeed, different methods sometimes produced large differences in H for the same individual (±0.3). Such large differences interfered with conclusions about whether a drawing sequence was persistent or anti-persistent because in some cases H could be both above and below 0.5, depending on the method.

By applying new methods of analysis, it will be possible to progressively grasp the ontogeny of drawing behaviour. The results of a single study are not sufficient to identify the development of a behaviour as complex as drawing. Only a grouping of clues, each one characterizing one aspect of the behaviour, could make it possible to comprehend the whole. This work on temporal fractal analysis provides one piece that was previously missing and completes our previous research on spatial fractal analysis of drawings. Such works seem promising to better understand the ontogeny of drawing behaviour and, by extending this type of analysis to other species, notably great apes, we could learn more about its evolutionary history.

## Acknowledgement

We thank Jean-Louis Deneubourg for his help on analyses. We are grateful to the school director and the teachers who gave us the opportunity to collect a large number of children’s drawings. We would like to warmly thank all the participants and parents of all the children who participated with enthusiasm to the study.

This project has received financial support from the CNRS through the MITI interdisciplinary programs.

## Author contributions

Conceptualization, B.B., L.M., C.S.; Methodology, B.B. and L.M.; Investigation, B.B., L.M. and C.S..; Writing – Original Draft, B.B. and L.M.; Writing – Review & Editing, C.S., A.M, X.M. and M.P.; Funding Acquisition, C.S. and M.P.; Resources, J.H.; Data Curation, B.B. and L.M., Supervision, C.S. and M.P.

## Resource availability

### Lead contact

Further information and requests for resources and reagents should be directed to and will be fulfilled by the Lead Contact, Lison Martinet

### Data and code availability

The dataset generated during this study are available at Zenodo : https://zenodo.org/record/5293436#.YStFntMzaYV

### Declaration of Interests

The authors declare no competing interests.

## Supplementary Materials

**Figure 1.**
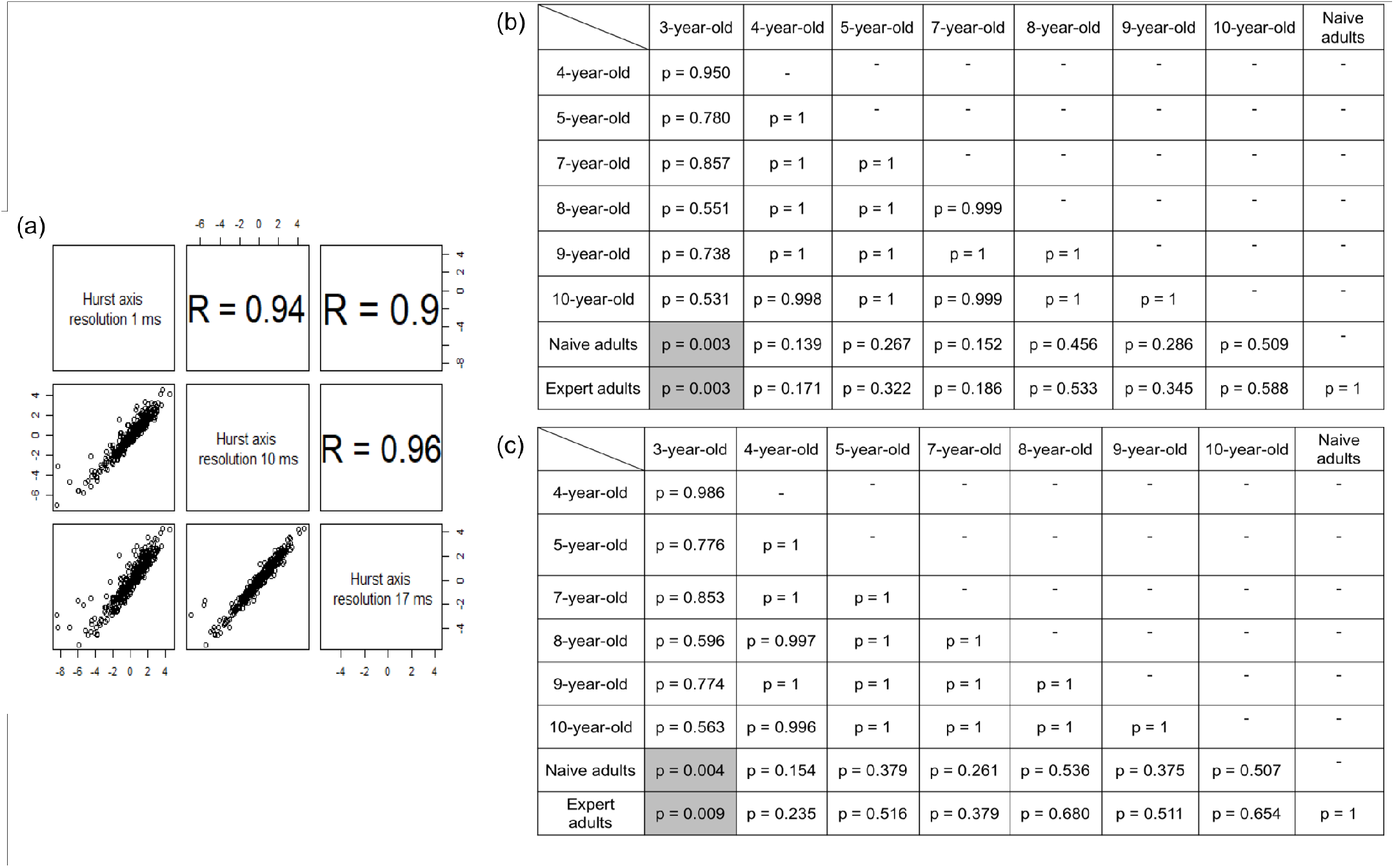
Analyses of different resolutions for the construction of the time series. (a) Pearson correlation matrix indicated the correlation between the Hurst axis values obtained with 3 different resolution in the time series (1ms being the one used in the manuscript). (b) Pairwise comparisons of groups obtained from the selected model (GLMM, *df* = 8, χ^2^ = 25.811, = 0.001) with a time resolution of 10 milliseconds. (c) Pairwise comparisons of groups obtained from the selected model (GLMM, *df* = 8, χ^2^ = 23.23, p = 0.003) with a time resolution of 17 milliseconds. The different resolutions give the same results with the 3-year-olds showing a higher value along the Hurst axis than adults, both, naïve and expert.

### Box 1.

Explanation of simulations showing the robustness of fractal estimates to random removals of segments within the sequence.

To understand the impact of removing a period of time in the sequences, and to ensure this will lead to consistent estimates, 50 drawings were randomly sampled. A chunk of 30 seconds was then randomly removed from the whole temporal sequence for each drawing, and the remaining parts were spliced. The 4 estimates were calculated on the first 50 seconds of the shortened sequences of drawings. The correlation of these estimates and the corresponding estimates on the first 50 seconds of the non-modified sequences were calculated. The correlations were strong (DFA: 97%, SWV: 96%, HAV: 92%, lowPSD: 94%), showing the consistency of this methodology and its robustness to missing segments of data.

**Figure 2.**
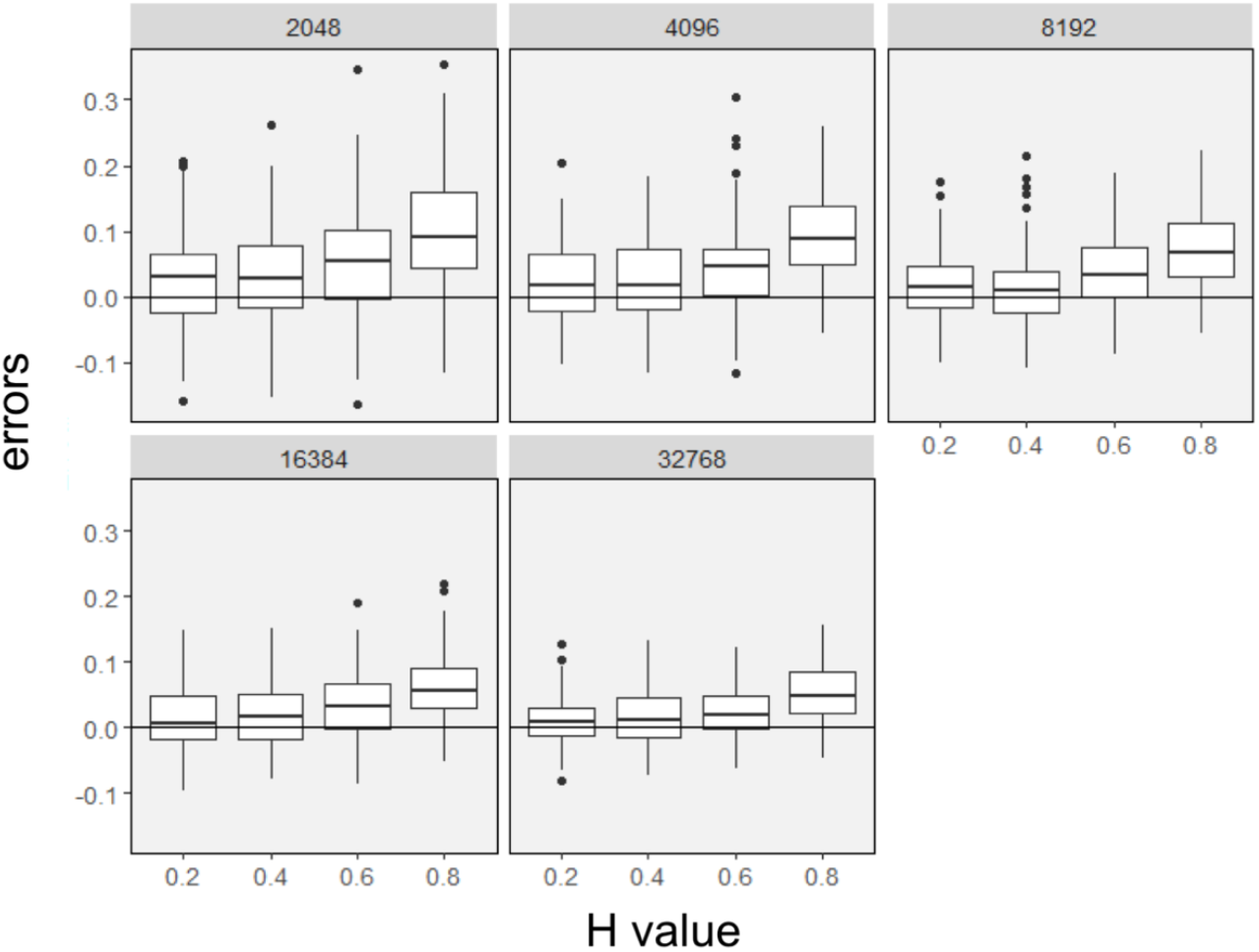
Estimation errors of the estimates of Monte Carlo simulations with the HAV method by varying H and the length of the time series.

**Figure 3.**
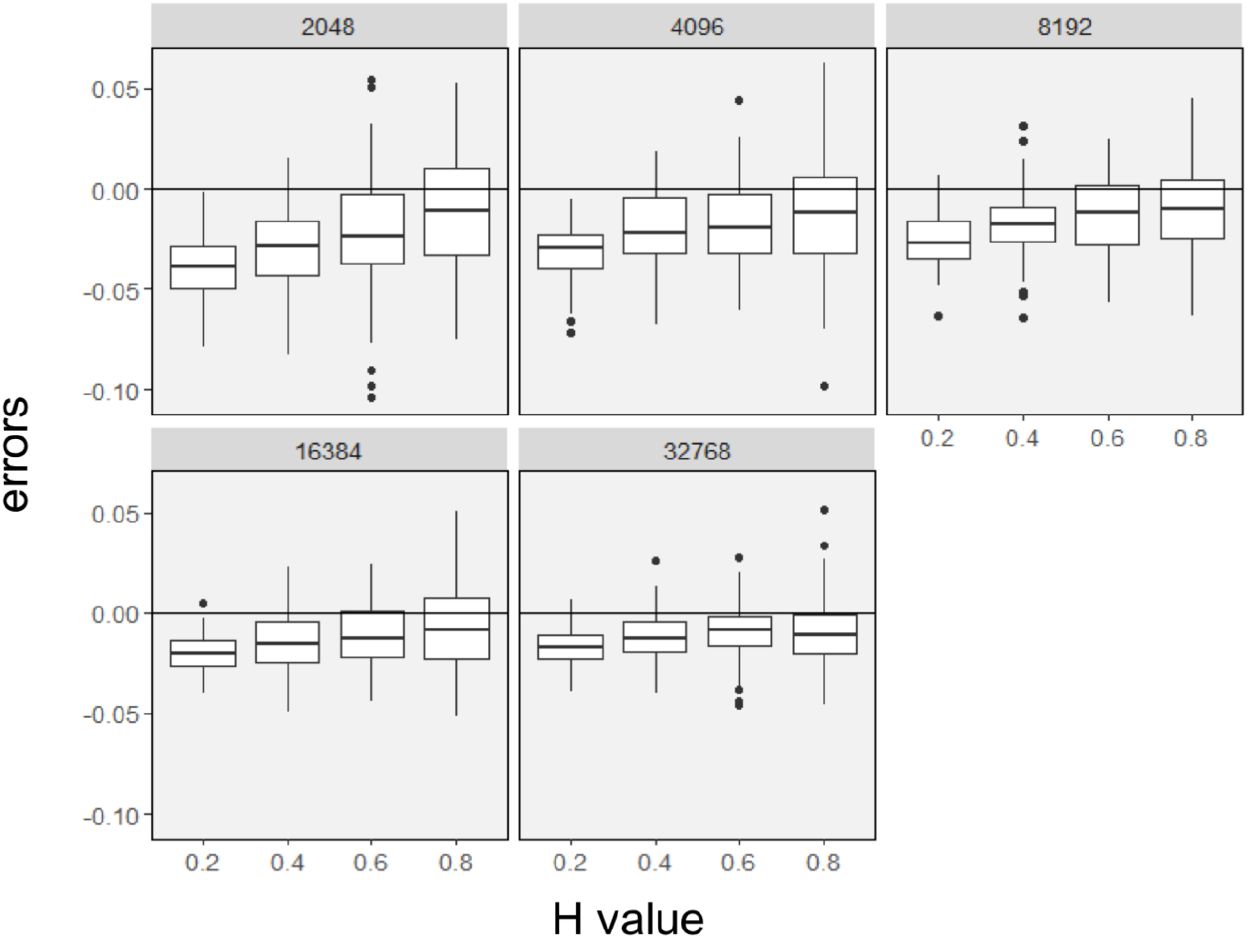
Estimation errors of Monte Carlo simulations with the SWV method by varying H and the length of the time series.

**Figure 4.**
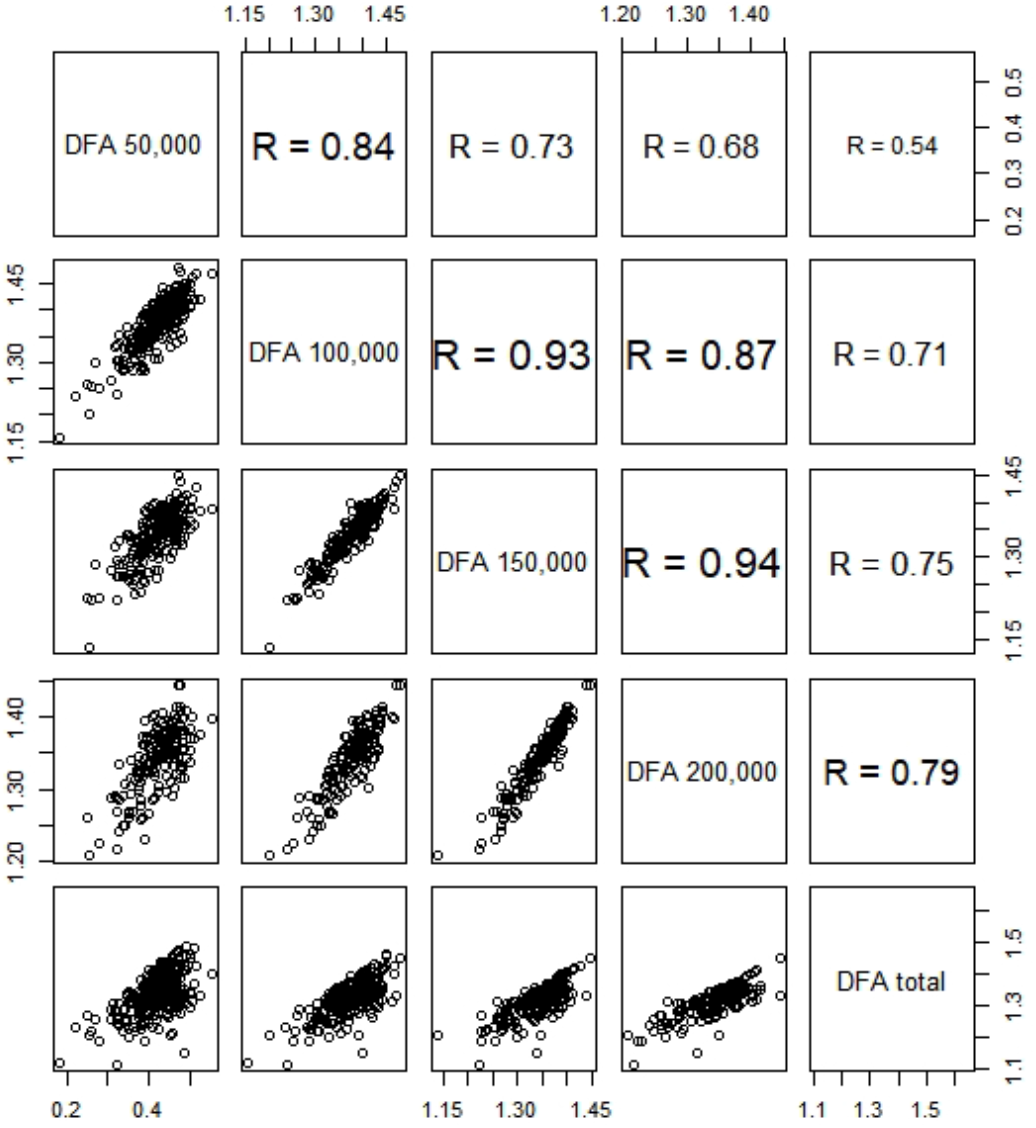
Correlations between DFA coefficients based on the first 50,000 points, 100,000 points, 150,000 points and 200,000 points and the total length of the times series.

**Figure 5.**
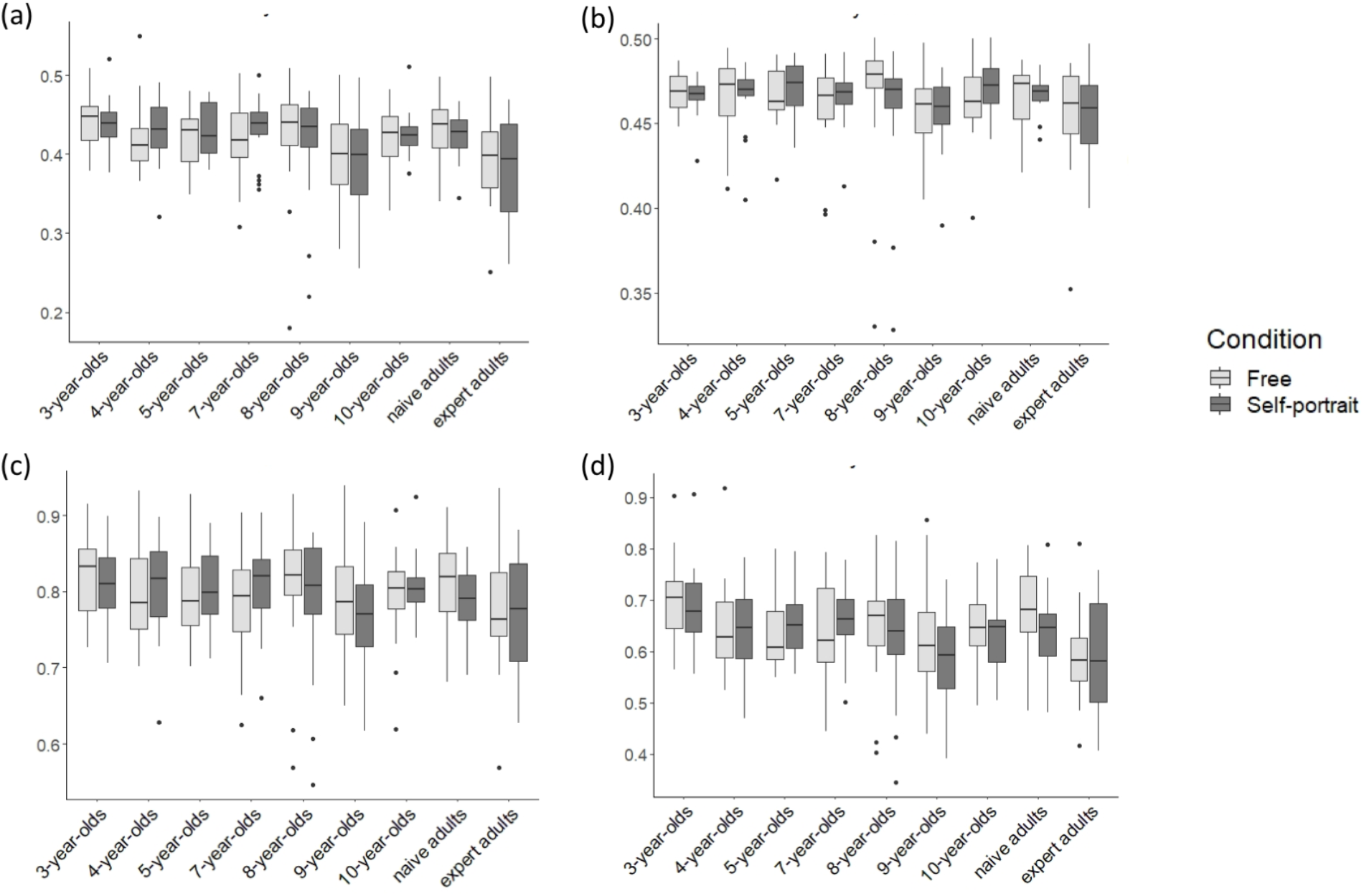
Boxplots of the Hurst estimates calculated with four different methods. (a) Hurst exponent estimated with (a) the Detrended Fluctuation Analysis (DFA), (b) the Power Spectral Density analysis (^low^PSD), (c) the Scaled Windowed Variance (SWV) and (d) the Hurst Absolute Value method (HAV).

**Figure 6.**
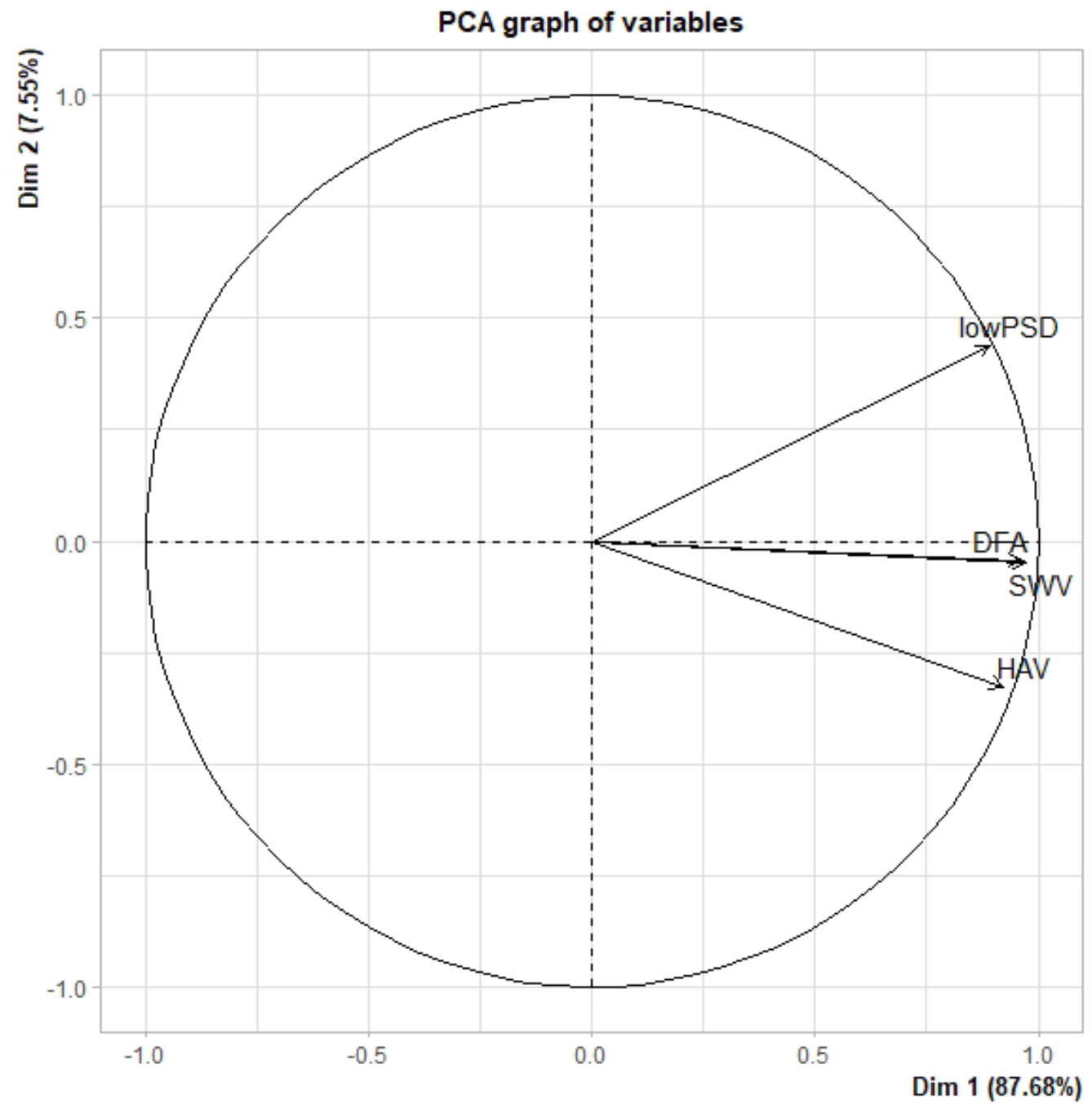
Correlation graph of the variables.

**Table 1.**
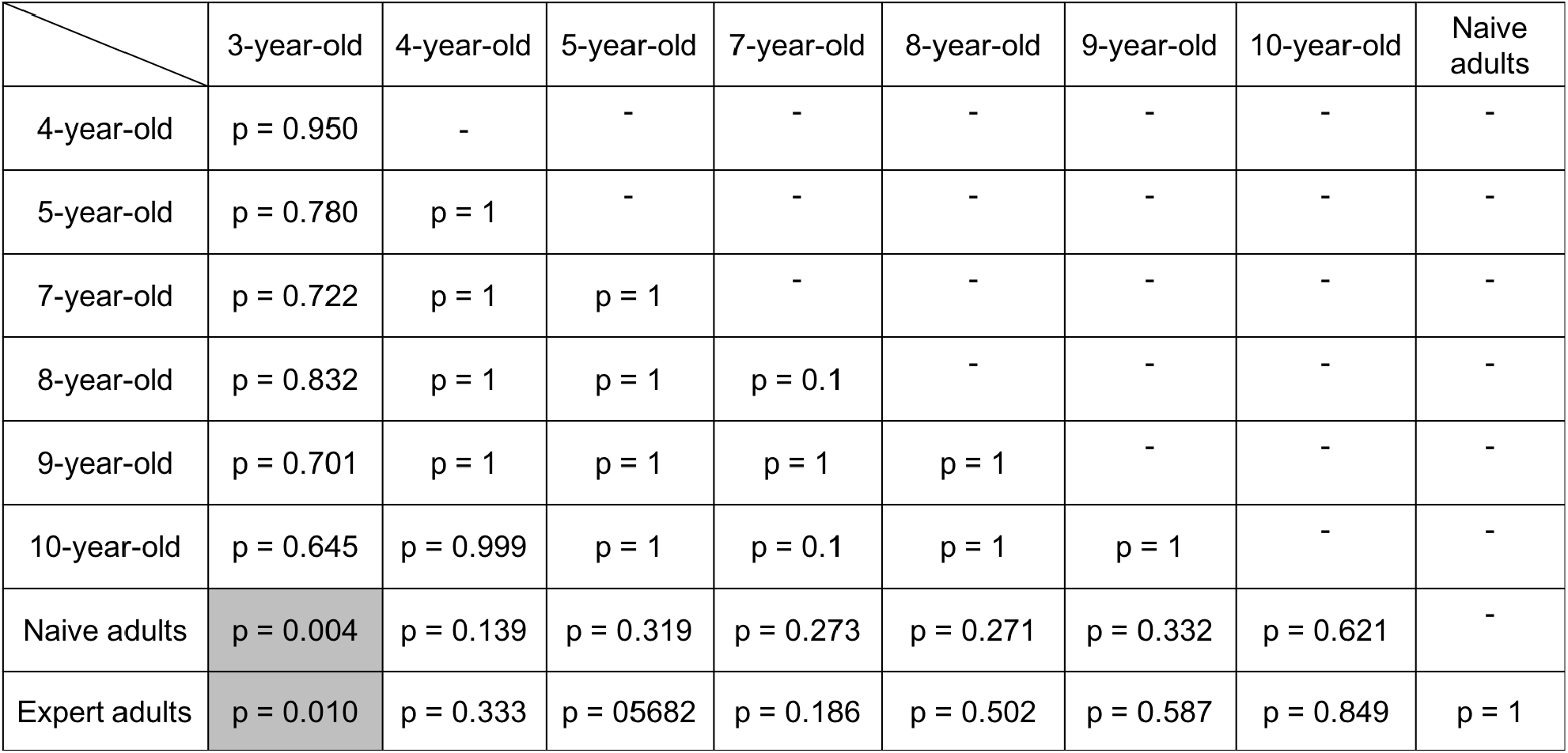
Pairwise comparisons of groups obtained from the selected model (GLMM, *df* = 8, χ^2^ = 22.842, p = 0.003) made by averaging the estimates of *H*.

## Notes

https://zenodo.org/record/5293436#.YStJkNMzaYW

### Competing Interest Statement

The authors have declared no competing interest.

